# Tuning spherical cells into kinking helices in wall-less bacteria

**DOI:** 10.1101/2021.11.16.467908

**Authors:** Carole Lartigue, Bastien Lambert, Fabien Rideau, Marion Decossas, Mélanie Hillion, Jean-Paul Douliez, Julie Hardouin, Olivier Lambert, Alain Blanchard, Laure Béven

## Abstract

In bacteria, cell shape is determined and maintained through a complex interplay between the peptidoglycan cell wall and cytoplasmic filaments made of polymerized MreB. *Spiroplasma* species, members of the Mollicutes class, challenge this general understanding because they are characterized by a helical cell shape and motility without a cell wall. This specificity is thought to rely on five MreB isoforms and a specific fibril protein. In this study, combinations of these five MreBs and of the fibril from *Spiroplasma citri* were expressed in another Mollicutes, *Mycoplasma capricolum. Mycoplasma* cells that were initially pleomorphic, mostly spherical, turned into helices when MreBs and fibrils were expressed in this heterologous host. The fibril protein was essential neither for helicity nor for cell movements. The isoform MreB5 had a special role as it was sufficient to confer helicity and motility to the mycoplasma cells. Cryo-electron microscopy confirmed the association of MreBs and fibril-based cytoskeleton with the plasma membrane, suggesting a direct effect on the membrane curvature. Finally, the heterologous expression of these proteins, MreBs and fibril, made it possible to reproduce the kink-like motility of spiroplasmas without providing the ability of cell movement in liquid broth. We suggest that other *Spiroplasma* components, not yet identified, are required for swimming, a hypothesis that could be evaluated in future studies using the same model.

## Introduction

Maintenance and dynamic reconfiguration of cell shape represent a selective value for bacteria for both primary and secondary cellular processes, in particular for nutrient acquisition, division, capacity to escape from predators, biofilm formation, and motility (Young, 2006). Therefore natural evolution has led most bacterial species to adopt one or a limited number of morphologies among many more or less complex possibilities, depending on their way of life and their ecological niche (Young, 2007). The main determinant of bacterial cell shape is the peptidoglycan layer surrounding the plasma membrane and forming the cell wall (Egan et al., 2020). The morphological transition of cells into spheres upon inhibition of cell wall synthesis in rod-shaped bacteria demonstrates the essential role of this structure in the maintenance of an elongated bacterial morphology (Claessen and Errington, 2019). In most rod-shaped bacteria, short internal filaments made up of actin-like proteins called MreBs guide the synthesis machinery of the cell wall to ensure cell elongation as deposition and crosslinking of new peptidoglycan units progress (Shi et al., 2018).

Along with cocci and rods, helical or corkscrew morphologies are major shapes adopted by phylogenetically distant bacteria including *Helicobacter pylori*, spirochetes and spiroplasmas. In *H. pylori*, the helical shape of the cell body can significantly contribute to propulsive thrust, (Constantino et al., 2016), or to pathogenicity (Salama et al., 2013). In this species, the cell wall is differentially synthesized based on the curvature of the cell body, with two proteins MreB and CcmA defining the appropriate areas where synthesis is enhanced (Taylor et al., 2020). Helical or wave-like morphologies and motility in spirochetes are primarily determined by the periplasmic flagella, the cell wall and cytoplasmic MreB (Nakamura, 2020). Spiroplasmas represent a group of helical bacteria apart. Indeed, spiroplasmas belong to the class Mollicutes, characterized by the lack of a peptidoglycan-based cell wall (Whitcomb, 1980). Most *Spiroplasma* species are pathogens or endosymbionts of arthropods and plants (Regassa, 2006). With the sole exception of the strain S*piroplasma citri* ASP-1, all natural *Spiroplasma* isolates described to date are helical and motile (Harne et al., 2020b) suggesting a selective value of this specific shape and unique motility. Thus, spiroplasmas control their helicity without a peptidoglycan layer, the major determinant of bacterial shape in walled bacteria. In addition, these bacteria are motile in the absence of external appendages such as flagella or pili that allow the motility of the vast majority of bacteria (Nakamura and Minamino, 2019). Recently, studies aiming at elucidating the mechanisms of shape maintenance and motility in *Spiroplasma* are increasing. Indeed, *Spiroplasma* appears to be a particularly attractive model for the identification of shape and motility determining factors in a wall-less organism (Harne et al., 2020b).

*Spiroplasma* cells are polarized, with a tapered (also called tip) and a rounded end (Garnier et al., 1984). The discovery of an internal cytoskeleton composed of the protein fibril (Fib) unique to *Spiroplasma* (Williamson, 1974) came closely after the discovery of these bacteria (Davis et al., 1972).The main cytoskeleton structure corresponds to a monolayered, flattened ribbon positioned along the shortest helical path (Trachtenberg and Gilad, 2001). Microscopic observations of cryofixed freeze-substituted preparations combined with tomographic reconstruction confirmed the overall organization and highlighted the membrane association of the internal protein ribbon made of both actin-like MreBs and fibril (Trachtenberg et al., 2008).The internal cytoskeleton also comprises a dumbbell-shaped structure at one cell pole (tapered-end) likely involved in cell polarization (Liu et al., 2017). The motility of *Spiroplasma* is due to a helicity change, which is initiated at one of the two ends of the cell and introduces a “kink” in the cell whose propagation is responsible for the movement of the bacteria (Shaevitz et al., 2005). On the cytoplasmic side, the complex dumbbell-shaped structure could be responsible for the generation of the initial twist which is then propagated along the cell (Sasajima and Miyata, 2021).

The presence of at least 5 *mreB* paralogs in the near-minimal genome of *Spiroplasma* species (Ku et al., 2014), raises some questions regarding the selective benefit provided by the different isoforms during evolution. In the non-helical strain *S. citri* ASP-1, the loss of helicity and motility is due to a nonsense mutation within the sequence encoding MreB5 that cannot be functionally compensated by any of the other 4 isoforms present in this species (Harne et al., 2020a).Thus MreB5 was identified as a major determinant of cell helicity in *S. citri* (Harne et al., 2020a).These *in vivo* results, coupled with differences in polymerization and depolymerization dynamics between isoforms (Masson et al., 2021; Pande et al., 2021; Takahashi et al., 2021), strengthen the hypothesis of functional differentiation between some MreB paralogs.

Despite the qualitative compositional information, structure and mechanisms of the *Spiroplasma* internal cytoskeleton remain unclear. Recently, based on the structure of the Fib filament, a molecular mechanism, in which fibril is responsible for cell helicity and for its shift has been proposed. Following this model, the length changes in MreB polymers would generate the helicity shift (Sasajima and Miyata, 2021). The construction of *Spiroplasma* mutants expressing combinatorial sets of MreBs with or without fibril could help validating this motility model and identifying the different roles of MreBs. However, although different tools have been developed to modify *Spiroplasma* genome, it remains difficult to obtain conditional gene knockdowns, especially for paralogs (Harne et al., 2020b). To circumvent these limitations, one approach is to perform heterologous expression experiments of *Spiroplasma* cytoskeleton proteins within a phylogenetically-related *Mycoplasma* species also lacking a cell wall. The choice of *Mycoplasma capricolum* subsp. *capricolum* (*Mcap*) is justified as it belongs to the Spiroplasma phylogenetic group of the Mollicutes class, and it is amenable to genome engineering including using methods derived from synthetic biology (Labroussaa et al., 2016; Lartigue et al., 2009). *Mcap* cells are coccoid and their genome does not encode any fibril or MreB protein. In addition, *Mcap* and *Spiroplasma* membranes have a similar lipid composition (Davis et al., 1985; Rottem, 1980), an essential point when considering that cytoskeleton elements were found to be closely associated with the membrane (Trachtenberg et al., 2008).

Here, we investigated whether the reconstruction of *Spiroplasma* cell structure in *Mcap* was possible. We then took advantage of this model to obtain clues on the minimal requirements for helical shape and kinking motility in Mollicutes, by comparing the effect of the insertion of different combinations of *S. citri* genes in *Mcap* genome on morphology, motility and formation of internal cytoskeleton elements.

## Results

### Expression of Spiroplasma cytoskeleton proteins in Mcap confers helicity and kinking motility

Recombinant strains of *M. capricolum* subsp. *capricolum* (*Mcap* hereafter) expressing *mreB* and fibril-encoding genes from *S. citri* were obtained after transformation with transposons (Fig. S1). As a control, *Mcap* cells were transformed with the transposon vector alone without additional genes (*Mcap*^*control*^). The morphology of the recombinant cells was initially analyzed by darkfield microscopy. Control cells were pleomorphic as expected for *Mcap, i*.*e*. they were mostly short-rod shaped and coccoid, a few elongated ones (up to 8 microns) but none with a helical morphology. In contrast, with the recombinants resulting from transformation with the transposon carrying the combination of *mreB1-5* and fibril genes and named *Mcap*^*mreB1-5-fib*^, the cell morphology was heterogeneous and included helices, long, straight and soft filamentous cells, eventually branched (up to 80 microns), but also partially helical cells, and coccoid cells (Fig. 1A). Helical pitch of the cells, corresponding to the distance between two equivalent points separated by a single turn on the helix, measured parallel to the cell axis and determined from darkfield microscopy images (insert, Fig. 1B), was 1.83+/-1.55 µm in average, a value significantly different from 0.74+/-0.3 µm determined for *S. citri* (Fig. 1B). Helical cells exhibited bending and kinking movements, mimicking those seen with *S. citri* (Fig.1C and Supplemental movie 1). However, kinks were initiated more irregularly than in *S. citri* and most of them did not spread down the entire length of the cell. Kinks locally travelled with a similar mean velocity V_kink_ in *S. citri* (V_kink_=12.0 ± 2 µm/s) and in *Mcap*^*mreB1-5-fib*^ (V_kink_=10.8 ± 2 µm/s). Motility with kink propagation was observed for helical pitches between 0.6 and 1.5 µm. Movements were also observed for non-helical filaments, with the propagation of a thrill along the cell body that induced bending of the elongated filaments.

**Figure 1.**
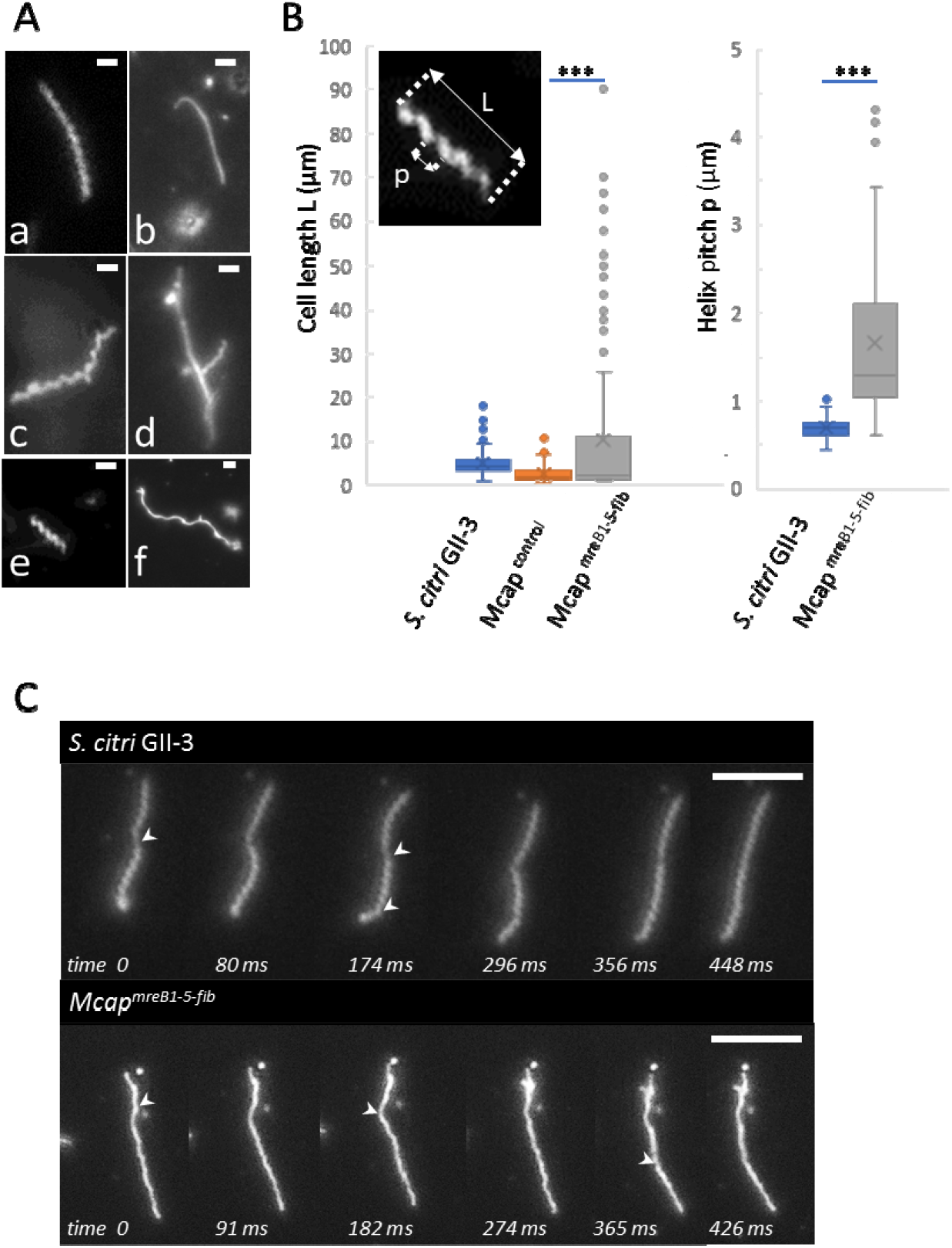
Morphology and motility of *Spiroplasma citri* GII-3 and *Mcap* transformants, in the genome of which *mreB* and fib genes of *S. citri* have been inserted. (A) Darkfield microscopy images showing representative morphologies of *Mcap*^*mreB1-5-fib*^ cells: Soft, long helical (a) and non-helical (b) cells; a branched, rigid helical cell (c); a straight, branched filament showing helical or straight lateral extensions (d); a short helical cell (e); and a long wavy cell (f). Scale bar: 2 µm. (B) Box plot display of the cell length and helical pitch in *S. citri, Mcap*^*control*^ transformed with the empty vector and *Mcap*^*mreB1-5-fib*^ cell populations measured using darkfield microscopy images. Top left insert represents the helical pitch p and the length L of a representative helical cell. Helical pitch measurements were restricted to helical cells. Data obtained with the different cell populations were compared with a two-tailed Student’s t-test, ***indicates a significant difference between populations with p<0.001. (C) Time-lapse images showing the kink-based cell movements in *S. citri* (top) and *Mcap*^*mreB1-5-fib*^ (bottom). White arrows point at kinks. Note the helicity shift upon propagation of the kink along the cell body. Scale bar: 5 µm.

The kink-based cell movements did not provide a directional motility to helical *Mcap*^*mreB1-5-fib*^ cells in liquid SP5. Since the medium viscosity minimizes Brownian motion and favors *S. citri* motility (Boudet et al., 2018), 0.2 to 1% methylcellulose was added to the medium but this additive did not improve the translational movement of the *Mcap*^*mreB1-5-fib*^ recombinant (data not shown). Also, while *S. citri* grown on agar-containing plates form satellite colonies due to the migration of single cells away from the mother colony, no satellites were observed with the *Mcap*^*mreB1-5-fib*^ recombinant after growth on nutrient medium containing 0.8% noble agar (Fig. S2), suggesting that the propulsive force conferred by the addition of *S. citri* cytoskeleton proteins is not sufficient to efficiently move the cells in one direction under the conditions tested.

Expression of MreBs and Fib in *Mcap* had not only a significant effect on cell morphology but also on cell division. Indeed, the presence of long filaments is likely the result of a defective septation during the cell division process. Elongation of the cells that were not correctly separated resulted in branching (Fig. 1A). Impairment in cell division was also correlated with a particular aspect of the colonies grown on SP4-agar medium: while the control transformants gave typical fried-egg colonies, *Mcap*^*mreB1-5-fib*^ recombinant growth resulted in more brownish and granular colonies, some with irregular contours (Fig. S2). Such colony morphology could be associated with division arrest for some cells.

### Fibril is essential neither for helicity nor for movements, but favors propagation of deformations at the membrane level along the cell body in Mcap

We took advantage of the *Mcap* heterologous system to assess whether or not helicity and motility could be conferred to the cells when *fib* only or *mreB1-5* only were added to the genome of recombinants (Fig. S3). To avoid the possible impact of the localization of the added genes in the *Mcap* genome (Table S1), only features common to all clones tested are described below.

*Mcap*^*fib*^ recombinants were characterized by a short-rod shape for most cells, but also by the presence of a few short helices having a helical pitch of 1.06+/-0.28 µm (Fig. 2). In the cell population of *Mcap*^*mreB1-5*^ recombinants, short and long helices were also observed, with a mean pitch of 1.72+/-0.67 µm, not significantly different from the pitch of *Mcap*^*mreB1-5+fib*^ transformants (1.83+/-1.55 µm). Thus, MreBs were sufficient to confer helicity to *Mcap* cells. It is noticeable that for both *Mcap*^*fib*^ and *Mcap*^*mreB1-5*^ transformants, the range of helical pitch lengths overlaps the length of that of *S. citri*, indicating that the *Spiroplasma* shape can be found with either of the constructions. Defects in septation were observed for most *Mcap*^*mreB1-5*^ cells, with the formation of long, branched filaments. One possible reason is that one or several MreBs may interfere with the division process. Also, some cell bodies showed certain flexibility, while others were characterized by a significant stiffness, possibly due to differences in amounts of cytoskeleton proteins. These problems in division were associated with the formation of brownish, irregular colonies on agar plates smaller than those observed with the control cells, *i*.*e. Mcap* transformed with the empty vector (Fig. S2).

**Figure 2.**
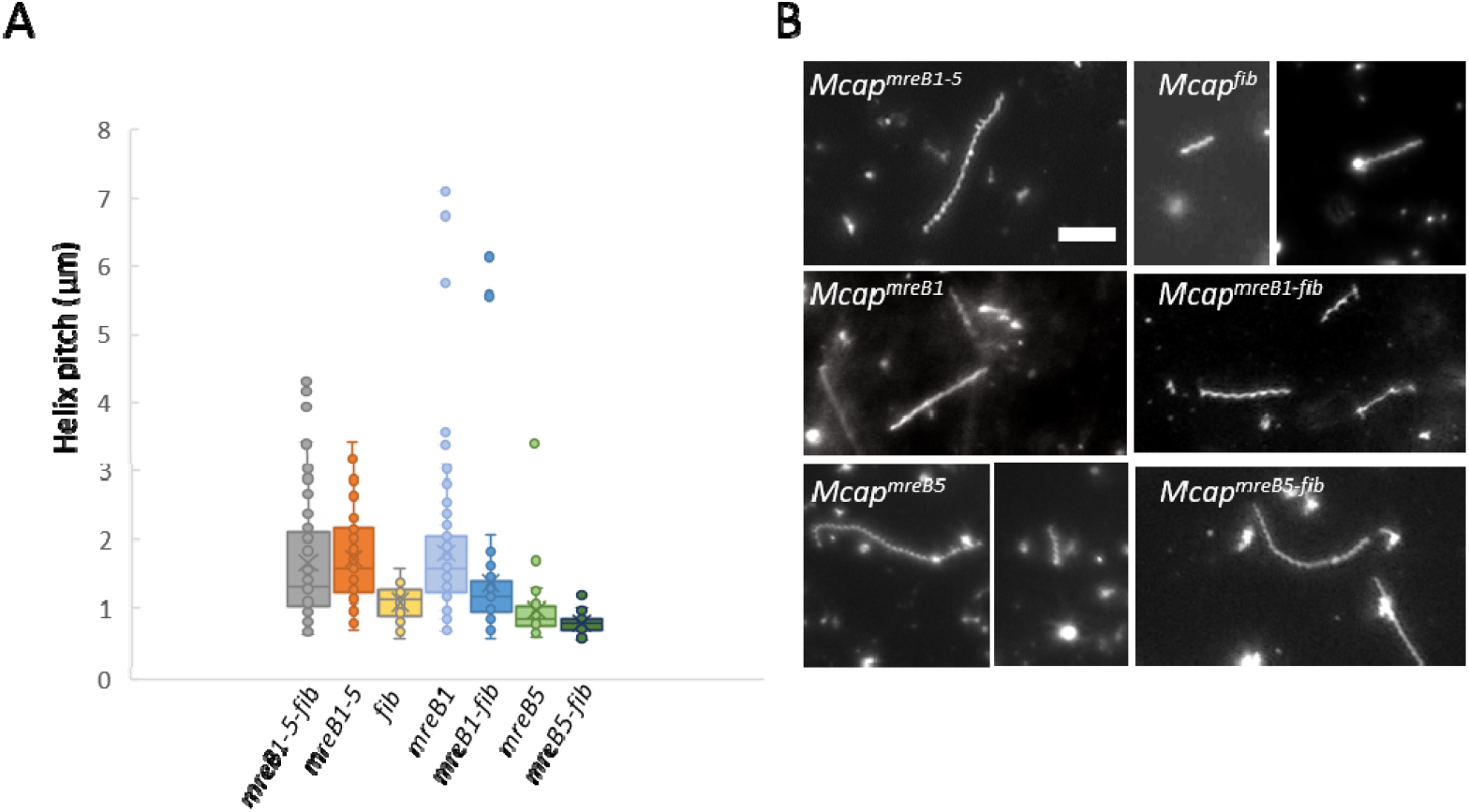
Helicity of *Mcap* transformants bearing different combinations of *mreB* and *fib* genes observed using darkfield microscopy. (A) Box plot display of the helical pitch in *Mcap* transformants. (B) Representative helical cells observed in *Mcap* transformed with the different plasmid constructs. Scale bar: 5 µm.

For both *Mcap*^*mreB1-5*^ and *Mcap*^*fib*^ recombinants, cell movements was observed for some helices with a helical pitch between 0.7 and 1.5 µm. Addition of both gene sets allowed the propagation of membrane deformations along the cell body, and was responsible for a change in helicity (Fig. 3). Thus, MreBs only or Fib only give *Mcap* the possibility of mimicking *Spiroplasma* membrane deformations responsible for its motility. These movements were however not sufficient to move the cells in one direction in liquid (Supplemental movie 2) or semi-viscous medium (data not shown). Motile helical cells were all of a short length. In addition, for *Mcap*^*fib*^ transformants, helical cells transiently lose their helicity during movements (Fig. 3).

**Figure 3.**
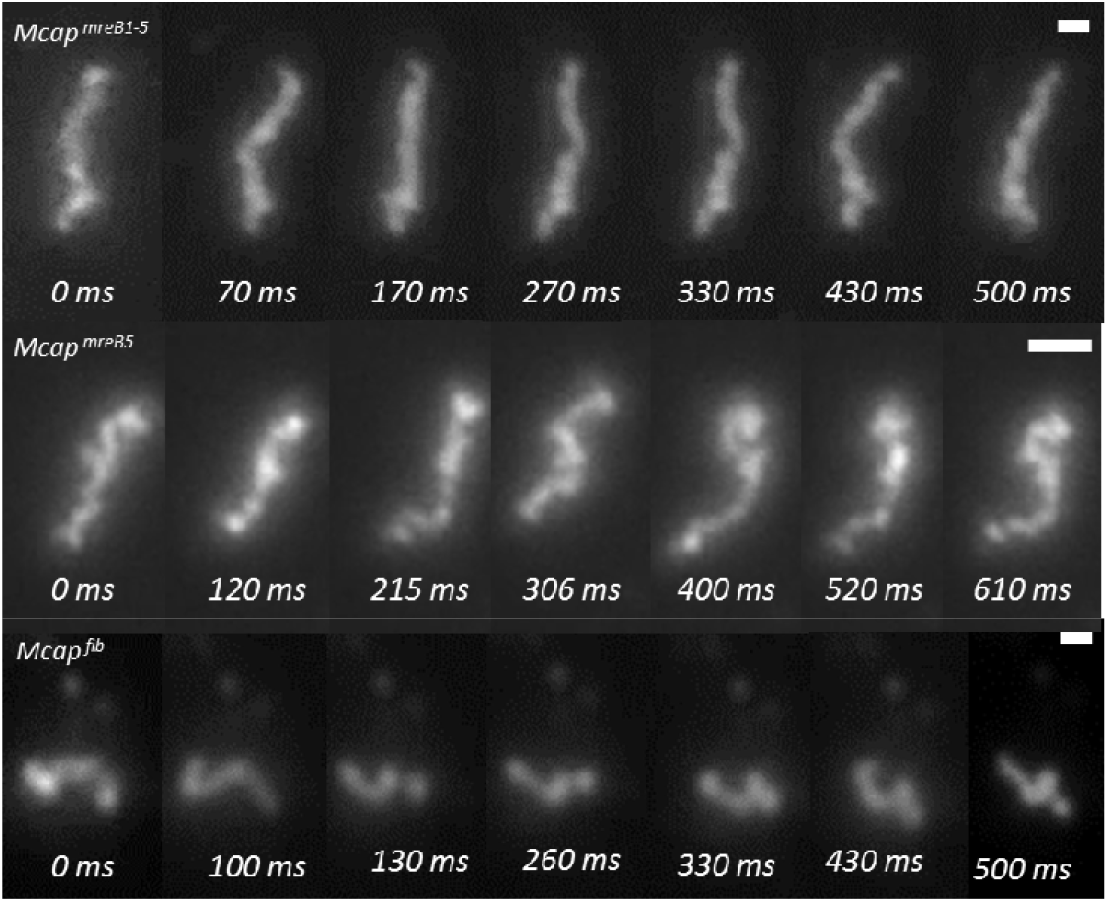
Cell movements in *Mcap* transformants. Darkfield microscopy time-lapse images showing helicity changes due to the propagation of membrane deformations in *Mcap* transformants bearing all *mreB* genes (top), *mreB5* only (middle) or *fib* only (bottom). Note that *Mcap* transformants having *mreB5* only or *fib* only lose their helicity upon propagation of the kink-like membrane deformation. Scale bar: 1 µm.

After 3 to 5 passages in liquid medium, helicity and motility were lost for the vast majority of the cells. A difference in motility between *Mcap*^*mreB1-5-fib*^ and *Mcap*^*mreB1-5*^ transformants was noticed. A large number of long filaments, essentially rigid with only a few, more flexible areas at their surface showed the propagation of a thrill along the cell body in both cell populations. However, only in the case of *Mcap*^*mreB1-5-fib*^, a bending of the rigid zone following a flexible point was observed. Taken together these results suggest that fibril is essential neither for helicity nor for motility, but likely increases the efficiency of propagation of the membrane deformation (kink) along the cell body. In addition, Fib can confer helicity and motility to short helices by itself.

### A single MreB is sufficient to confer helicity and to initiate kinks

Proteomics analyses were performed to check the expression of MreBs in *S. citri*, in one clone of *Mcap*^*mreB1-5*^ and two clones of *Mcap*^*mreB1-5+fib*^. For one *Mcap*^*mreB1-5+fib*^ clone, protein expression was also analyzed after 4 and 8 passages in axenic medium. Morphology of exponentially growing cells was checked before extraction of total proteins and LC-MS/MS analysis. MreB5 was the most expressed among MreBs in these transformants, as in *S. citri* (Table 1). Surprisingly, although all *mreB* genes were added in all clones, not all MreBs were detected by proteomics. Even more unexpected was the observation that the set of detected proteins was different from one clone to another. More generally, relative abundance of MreBs and Fib in *Mcap*^*mreB1-5+fib*^ did not mimic those in *S. citri*. Fib was detected at a significantly lower level in *Mcap*^*mreB1-5+fib*^, and MreB5 amount after 8 passages in SP5 tet5 was more than 20 times higher than after 4 passages. All clones having shown to be able to form motile helices, different combinations of MreBs were then able to confer helicity and motility. MreB1+MreB5, MreB4+MreB5, MreB1+MreB4+MreB5, and all MreBs were among the MreB combinations generating motile helices (Table 1). It should however be stressed out that the apparently lacking MreBs may be expressed at a level too low to be detected. Aiming at determining the minimal gene set required for helicity and/or motility, a transformant carrying only either *mreB*5 or *mreB1* was constructed (Fig. S4).

**Table 1.**
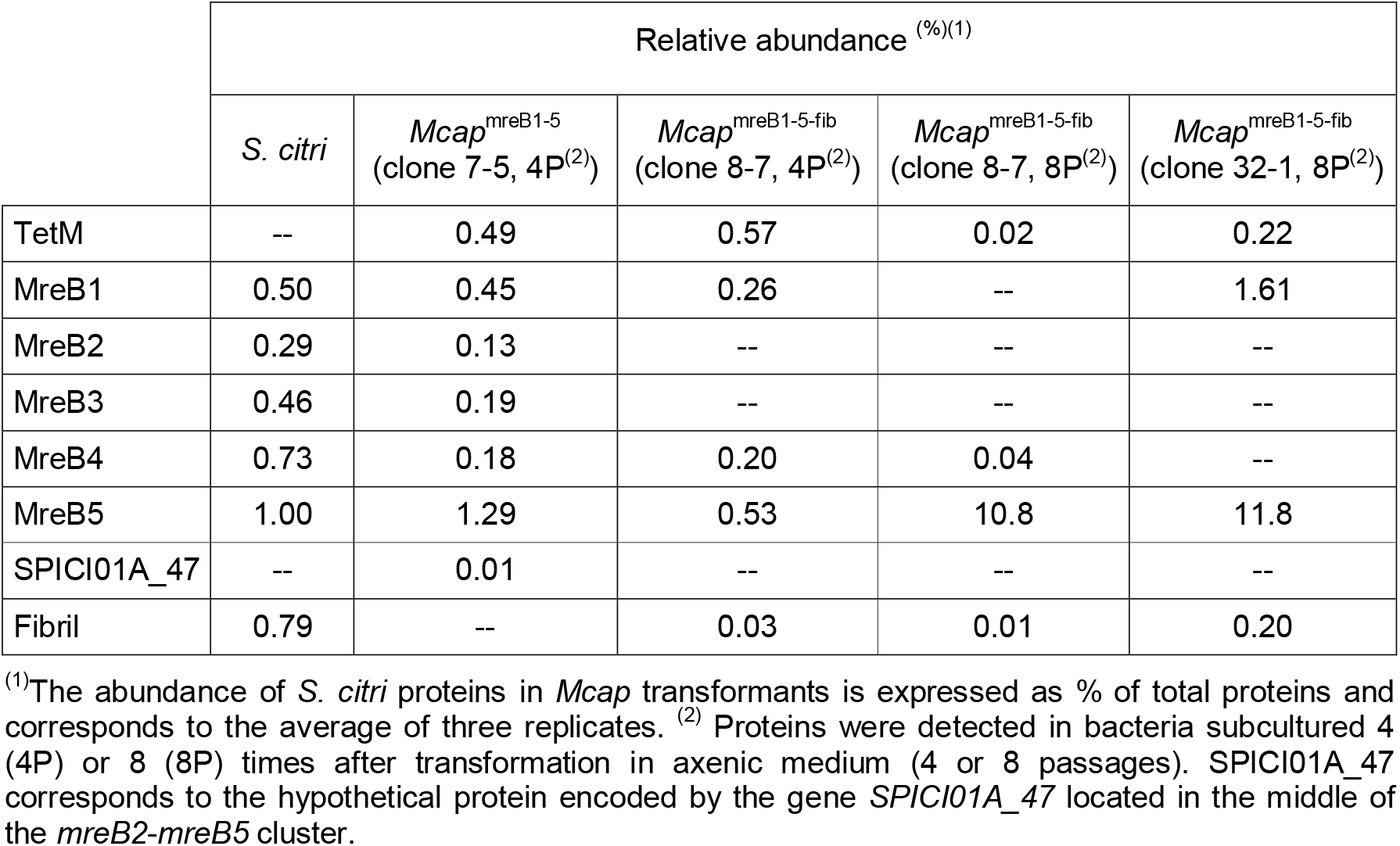
Heterologous expression of MreBs and Fib in *Mcap* transformants.

While *Mcap* transformation efficiencies with all the constructions (except for pMT85-PStetM-fib) were rather low (Table S2), the one with pMT85-PStetM-*mreB5* was even lower. Nevertheless, two clones could be recovered after three transformation assays. *Mcap*^*mreB5*^ population showed a large diversity of shapes from coccoid to filamentous ones, including helices with a helical pitch of 0.96+/-0.48 µm, which was not significantly different from those of *S. citri* helices. Short helices with a length inferior to 5 µm were also endowed with movements, associated with the propagation of membrane deformations along the cell body (Fig. 3). Although the change in helicity could be visualized (Supplemental movie 2), the cellular stiffness was not sufficient enough to conserve the helicity on the whole length during movements, which led to the transient formation of unwound and untangled filaments. Longer helices were not found to be motile. Subculturing of the *Mcap*^*mreB5*^ clones led to loss of helicity and motility.

When *mreB5* was co-expressed with *fib* (*Mcap*^*mreB5-fib*^) (Fig. S5), more stable helices were produced, and the helical pitch was 0.77+/-0.13 µm (Fig. 2). However kink-based motility was not observed with the gene combination *mreB5-fib*.

All these observations show that addition of *mre*B5 is sufficient to confer helicity to *Mcap* and triggers kink-like membrane deformations for short helices. The morphology of *Mcap*^*mreB5*^ was then compared to those of *Mcap*^*mreB1*^ carrying *mre*B1 gene alone (Fig. S4). The latter shows helices with a larger pitch (1.81+/-1.08 µm) (Fig. 2). Twitching movements were observed on flexible, non-helical cells, but kink-based motility of helices was not recorded with *Mcap*^*mreB1*^. Addition of *fib* gene together with *mre*B1 (Fig. S5) produced non-motile, stable helices with a mean pitch of 1.35+/-0.97 µm.

### Expression of MreBs and Fib was associated with formation of cytoskeleton structures interacting with the plasma membrane

CryoEM was used to assess whether the different gene combinations led to formation of stable cytoskeleton filaments in *Mcap*. Fig. 4A illustrates the tapered end of *S. citri* GII-3 and its internal cytoskeleton closely associated with the membrane at locations with negative curvature. Image of a typical control cell corresponding to *Mcap* transformed with the empty vector lacking any internal cytoskeleton structure is shown Fig. 4B. Upon adsorption on carbon grids and cryofixation prior to cell imaging, most helical *Mcap* recombinants lost their morphology. Nonetheless, the cells showed internal cytoskeleton elements. No cell polarization could be observed in *Mcap* recombinants whatever the gene combination.

**Figure 4.**
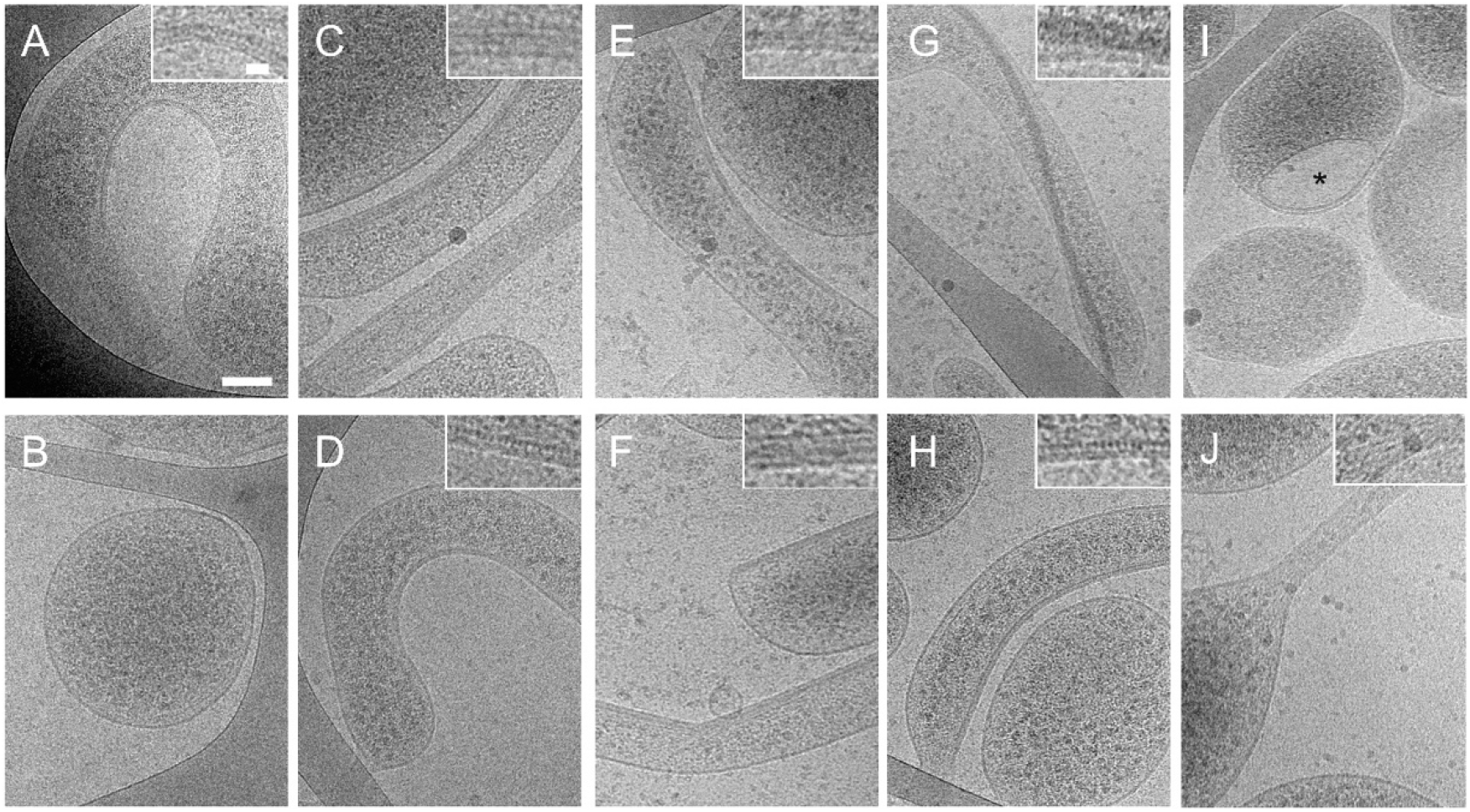
2D cryo-electron microscopy reveals the formation of cytoskeleton filaments in *Mcap* transformants bearing *S. citri* cytoskeleton genes. (A*) Spiroplasma citri* cytoskeleton fibers are localized in the cytoplasm next to negatively curved areas of the plasma membrane. Note the tapered (duckbill-shaped) tip of the cell; (B) *Mcap*^*control*^ cells transformed with the empty vector do not show any cytoplasmic cytoskeleton fibers; (C-H) Cytoskeleton fibers were imaged in *Mcap*^*mreB1-5-fib*^ (C), *Mcap*^*mreB1-5*^ (D), *Mcap*^*mreB1*^ (E), *Mcap*^*mreB1-fib*^ (F), *Mcap*^*mreB*5^ (G), *Mcap*^*mreB5-fib*^ (H) cells; (I) Internalized vesicle (indicated by *) observed in some *Mcap*^*mreB5*^ cells (J) Cell protrusions containing cytoskeleton fibers in *Mcap*^*fib*^ cells;. Scale bars: 100 nm and inset 20nm.

As explained above, addition of the complete gene set (*mre*B1-5 together with *fib*) in *Mcap* resulted in different expression profiles. Here the clone 8.7 (4P) expressing MreB1, MreB4, MreB5, and Fib was analyzed using cryoEM. Expression of these proteins induced the production of internal fibers positioned next to the membrane (Fig. 4C). The cytoskeleton could recover the whole inner side of the membrane in some cells, or appear only next to negatively curved membrane areas in others. The latter organization mimicked the *Spiroplasma* one, and differences observed between cells were likely due to differences from one cell to another in protein amounts. The internal cytoskeleton structure in *Mcap*^*mreB1- 5-fib*^ corresponded to a stack of filaments with a mean width ranging from 4 to 10 nm, and showing a regular striated pattern with a 5+/-1nm periodicity. Straight cells in which the cytoskeleton covers most of the membrane could correspond to cells showing a significant stiffness observed in cell suspensions as described above.

An internal structure was also observed in *Mcap*^*mreB1-5*^, *i*.*e*. in the absence of *fib* (Fig. 4D). The internal filament formed by one or several MreBs ran next to the membrane, in close interaction either with curved membrane areas, or with a larger membrane zone. The width of protofilaments ranged from 4 to 8 nm. The addition of *fib* together with at least one *mreb* triggered the formation of internal filaments associated with the membrane and showing the regular striated pattern described for *Mcap*^*mreB1-5-fib*^ (Fig. 4C, F, H).

*Mcap* recombinants expressing a single cytoskeleton protein (Fib or MreB1 or MreB5) showed also internal structures. Expression of MreB1 induced the formation of stable cytoskeleton fibers adjacent to the membrane (Fig. 4E). Filaments made of MreB1 (width ranging from 4 to 8 nm) conferred rigidity to cells when interacting with large membrane segments. These internal structures can recover the entire membrane inner side. In *Mcap*^*mreB5*^, MreB5 filament was clearly observed only in a single cell (Fig. 4G). MreB5 filament, having a width of up to 27 nm, crossed the whole cell body with localized interactions with the plasma membrane. In the absence of any MreB, expression of Fib allowed the formation of an internal cytoskeleton made of a stack of approximately 10 nm wide filaments, in particular in some tubular protrusions observed for some cells (Fig 4J). Interestingly, a large number of internalized membrane-bound structures were observed in *Mcap*^*mreB1-5*^, *Mcap*^*mreB1*^ and, even more frequently, in *Mcap*^*mreB5*^ (Fig 4I). This observation is in line with the great ability of MreB1 and MreB5 to induce membrane curvature at directed membrane areas, as an excessive curvature could lead to inverse membrane blebbing followed by detachment and internalization of a more or less spherical membrane-bound vesicle.

## Discussion

### Identification of the minimal requirements for conferring helicity and motility to Mcap

The *Spiroplasma* cytoskeleton was reconstituted in a mycoplasma that was initially pleomorphic and non-motile. Heterologous expression of MreBs and fibril proteins in *Mcap* cells changed the cell morphology from spherical to elongated cell bodies. In addition, it was sufficient to confer helicity and kinking motility to the mycoplasma cell. All *Spiroplasma* species have at least 5 copies of the *mreB* gene (Ku et al., 2014), which raises the question of the redundancy of functions between the different isoforms. Fib, MreB5 or MreB1 could generate cell helicity in *Mcap*, indicating that each of these proteins was able to induce the membrane curvature required for helicity in *Mcap*. The combination of Fib with MreB1 allowed the tightening of the helices, but the mean helical pitch remained higher than those of *S. citri* cells. On the contrary, MreB5 expression produced helices with a mean pitch similar to those found in *S. citri*. Hence each of these proteins represents a minimal requirement for helicity in a wall-less bacterium, and MreB5 the minimal one to mimic *Spiroplasma* helicity.

Regarding motility, the expression of MreB1 induced disorganized cellular movements, resembling tremors, but the observed membrane deformation did not propagate along the cell body, while MreB5 was able to confer kink-based motility to short helices. Therefore, a single MreB, such as MreB5, represents the minimal requirement to produce helical cells endowed with kink-like motility. However, the cytoskeleton resulting from the expression of a single MreB was not sufficient to allow the conservation of the cell length and helicity upon propagation of membrane deformation. Changes in cell length were likely responsible for loss of helix directionality. A similar phenotype was obtained when transforming *Mcap* with *fib* only. In *Spiroplasma*, fibril forms polymers whose length remains the same during motility (Sasajima and Miyata, 2021), ensuring the efficient conservation of the helical shape when the kink travels along the cell. In *Mcap*, the sole addition of *fib* did not provide the required helix stiffness. As discussed above, the co-expression of Fib with MreB5 allowed the stabilization of long helices, suggesting that a MreB could help in stabilizing the Fib-based cytoskeleton. However, in this configuration, the energy transfer to the membrane coupled by MreB5 and by Fib was not sufficient to induce membrane deformations propagating along the entire cell length. This could be due to the limited efficacy of the Fib-MreB5 combination in deforming a cell, which had become more rigid thanks to the presence of a Fib filament as compared to recombinants expressing a single *Spiroplasma* cytoskeleton protein. The expression of only MreB4 and MreB5, or of MreB1 and MreB5 together with *fib* in different *Mcap*^*mreB1-5-fib*^ clones was required to produce long kinking helices. This observation provides some clues in favor of the hypothesis of at least partially overlapping functions for MreB1 and MreB4 in *Spiroplasma*. The phylogenetic tree based on MreB sequences from 26 *Spiroplasma* species revealed that these proteins could be classified into 5 clusters (Harne et al., 2020a). However several MreBs, including *S. citri* MreB1, could not be clearly classified as MreB1 or MreB4, arguing in favor of our hypothesis of overlap of functions between these isoforms. The expression of MreB5 in all helical and motile *Mcap* transformants is in agreement with a major role played by this specific MreB in *Spiroplasma* helicity and motility. This result is consistent with the restoration of helicity and motility observed upon complementation with *mreB5* WT gene in *S. citri* ASP-1 (Harne et al., 2020a). It is noteworthy that the absence of a functional *mreB5* gene in the ASP-1 strain could not be compensated by any of the other 4 *mreB* genes, strongly suggesting a functional specialization between MreB5 and the other isoforms.

### Insights into the structure of *Spiroplasma* MreB filaments

*Spiroplasma* MreB3 and MreB5 were previously shown to polymerize *in vitro* (Harne et al., 2020a; Pande et al., 2021; Takahashi et al., 2021), and Masson et al. (2021) recently demonstrated that *Spiroplasma* MreBs expressed in *E. coli* could form a complex network supporting the hypothesis that the different isoforms participate to the production of a cytoskeleton in *Spiroplasma*. The present work not only indicates that *Spiroplasma* MreBs polymerize into stable filaments in a wall-less bacterium phylogenetically close to *Spiroplasma*, but also provides clues regarding the cytoskeleton organization. In *Mcap*, the width of MreB filaments could be as thin as 4 nm, similar to those of unidentified filaments previously observed in *S. citri* (Trachtenberg et al., 2003), which validates our heterologous system. In rod-shaped, walled bacteria, MreBs establish a direct interaction with the plasma membrane (Salje et al., 2011). The capacity of MreB5 to interact with lipid bilayers *in vitro* was previously demonstrated (Harne et al., 2020a). In *Mcap*, both MreB1 and MreB5 induced a plasma membrane curvature and formed filaments that were closely associated with the membrane. This ability to distort membranes is common to some other MreBs, as shown for TmMreB, an MreB from the thermophilic archaeon *Thermotoga maritima* (Salje et al., 2011). In some cells, expression of each of the MreBs in *Mcap* led to internalization of membrane bound vesicles. Interestingly, similar vesicles were observed in *E. coli* expressing TmMreB (Salje et al., 2011), strengthening the hypothesis that *Spiroplasma* MreB1, MreB5 and TmMreB share functional features. In the absence of any resistance provided by a peptidoglycan wall in Mollicutes, the curvature induced by *Spiroplasma* MreBs could lead to the formation of a helical cell. Although both MreB1 and MreB5 seem to directly interact with the membrane, the analyses of helical pitches in the different constructs lead us to put forward the hypothesis that MreB5 is the main determinant of the cell curvature allowing the correct localization of interactions between the internal ribbon and the plasma membrane. To induce the formation of helices having a regular pitch, *Spiroplasma* MreBs must interact with specific membrane partners. Considering that MreB5 interacts with liposomes (Harne et al., 2020a), its membrane partner in *Spiroplasma* and in *Mcap* may be of a lipidic nature. Since anionic phospholipids exclude assembled MreB in *E. coli* (Kawazura et al., 2017), MreB1 and B5 could preferentially interact with anionic phospholipids-depleted membrane areas in *Mcap* and in *Spiroplasma*. A heterogeneous distribution of phosphatidylglycerol and cardiolipin, major anionic phospholipids in *Mcap* and *Spiroplasma* membranes (Rottem, 1980), could occur and favor the interaction of *Spiroplasma* MreBs with membrane parts enriched in specific lipids. Of note, bundles of thick filaments were associated with rigid, straight cell morphology, indicating that helicity likely requires a thinner filament. Also, our cryoEM experiments indicate that MreB1 and B5 filaments could span the whole *Mcap* cell length over several micrometers. In most rod-shaped bacteria, the current prevalent hypothesis assumes that MreBs form short membrane-associated filaments (Shi et al., 2018) but their length is still debated (Errington, 2015). The length of MreB1 and B5 filaments in *Mcap* may differ from those of MreB filaments in *Spiroplasma*, as their expression level is probably not the same in the two species.

### Role of MreBs in motility

The demonstration of a role of MreB in bacterial motility has so far only been obtained in gliding motility of *Myxococcus xanthus* (Treuner-Lange et al., 2015). A role for MreB5 in *Spiroplasma* swimming was also suggested since the complementation of the non-helical, non-motile *S. citri* ASP-1 with *mreB5* from the helical and motile *S. citri* GII-3 restored not only helicity but also kinking motility (Harne et al., 2020a). Nonetheless, *Spiroplasma* kinking motility requires helicity. Therefore, motility restoration in *mreB5*-complemented ASP-1 could be due to helicity recovery and the role played by MreB5 in motility could then be indirect. Differential polymerization kinetic parameters of MreB isoforms measured *in vitro* led Sasajima and collaborators (Sasajima and Miyata, 2021) to propose a theoretical model for *Spiroplasma* swimming mechanism in which two MreBs would generate a force similar to that of a bimetallic strip. This force would then be transmitted to fibril polymers and result in change in handedness of helical fibril filaments. This model would fit our data showing that expression of two distinct MreBs allowed both helicity and kinking motility. However, a single MreB, MreB5, was also able to endow *Mcap* cells with movements resulting from the spread of a membrane deformation along the cell body. In most species, MreB filaments are highly dynamic polymers that align along the greatest principal membrane curvature (Hussain et al., 2018) and show circumferential motion following a path mostly perpendicular to the long axis of the cell (Morgenstein et al., 2015). In rod-shaped, walled bacteria, this motion is powered by the activity of the peptidoglycan synthesis machinery (Domìnguez-Escobar et al., 2011). Without peptidoglycan synthesis enzymatic activity in Mollicutes, assembly and dynamics of MreBs may drive the initiation and propagation of membrane deformations at the cell surface. The presence of different MreB isoforms in *Spiroplasma* could allow the generation of a cumulative force transmitted to the membrane.

### Role of fibril in *Spiroplasma* shape and motility

The present work also sheds light on the possible involvement of the fibril in the generation of helicity, its maintenance and motility. Fib filaments were observed in close interaction with *S. citri* plasma membrane in previous studies (Kürner et al., 2005; Trachtenberg et al., 2008). It was long thought that fibril was responsible for *Spiroplasma* motility by changing its length (Kürner et al., 2005). However recent studies indicated that the length of Fib polymers do not change during cell movement (Sasajima et al., 2021). The structure of Fib filaments has been studied by electronic microscopy: Fib filaments show an helicity with a pitch close to those of the *Spiroplasma* cell, leading to the conclusion that Fib was the determinant of helicity in *Spiroplasma* (Sasajima et al., 2021). Our results indicate that Fib is able to form membrane-interacting polymers in *Mcap* with a width similar to those observed in Fib isolated filaments (Trachtenberg et al., 2003; Sasajima and Miyata, 2021), and induce membrane curvature in the absence of any MreB. However, the helix constructed with Fib only is in a more relaxed form than those observed when MreB5 and Fib were co-expressed, suggesting that MreB5 could help positioning the Fib filament by interacting with both Fib and the membrane. This is in line with the ability of MreB5 to bind both fibril and liposomes *in vitro* (Harne et al., 2020a). Given its well-adapted helical structure and the rigidity it confers to cells, Fib could be a major determinant in helical shape maintenance. Regarding long helix generation, Fib had to be associated with MreB5 in *Mcap*, and probably in *Spiroplasma* also. Our work also provides some clues about the role played by Fib in motility. Fib allowed *Mcap* to form helices endowed with a motility very similar to those observed with MreB5. This is intriguing because Fib and MreB do not share any sequence similarity, and unlike MreBs, Fib lacks ATPase activity (Sasajima et al., 2021). In the most recently proposed model for *Spiroplasma* motility, MreBs transmit the force to the fibril filament which changes its handedness to generate the shift in the cell body helicity (Sasajima and Miyata, 2021). We propose that fibril is not only an essential structural component for transmission of the torque to the membrane, but also participates itself to the generation of the required force. Interestingly the genome of a few motile *Spiroplasma* species including *S. sabaudiense* (Abalain-Colloc et al., 1987) lacks the *fibril* gene (Chang et al., 2014), but contains more than 5 *mreB* genes (Harne et al., 2020b; Ku et al., 2014). It is therefore tempting to suggest an evolutionary convergence between MreB and fibril to ensure motility.

### Lack of an efficient swimming in *Mcap* recombinants

It is noteworthy that none of the *Mcap* recombinants exhibits translational motility in liquid medium. Among the hypotheses that can explain this observation is the lack of cell polarization in recombinants. It is reasonable to assume that a firm attachment of the internal cytoskeleton ribbon at the two poles of the cells is required to ensure translational movement. In *Mcap* recombinants, such attachment may be lacking. In addition, the *Spiroplasma* dumbbell-shaped core (Liu et al., 2017) was absent in *Mcap* transformants. We could show here that this structure was not required for kink generation and propagation, but it may be essential to generate an initial kink located at the extremity of the cell. Also, the tapered shape of the *Spiroplasma* tip may favor the penetration and propulsion in liquid media. Finally, the relative abundance of the proteins did not match those observed in *S. citri*. More specifically, Fib was detected at a low level, whereas the level of expression of MreB5 was deregulated, as indicated by its increase during subculturing in *Mcap*^*mreB1-5+fib*^. A proper stoichiometry of these proteins is likely to be required for optimal cell stiffness, stability of the helical structure in *Mcap* recombinants during the propagation of the kink, and efficient swimming.

## Conclusion

The present study demonstrates the importance of structure and organization of MreBs and fibril in determining helicity and motility in *Spiroplasma*. It provides some clues that leads to a structural model in which (i) the MreB5 filaments interact specifically with membrane areas enriched in yet unidentified lipids and induce membrane curvature, (ii) MreB5 determines the correct localization of MreB1 and fibril, which both participate to the membrane curvature, (iii) MreB1, MreB5 and Fib polymerize into filaments along the plasma membrane following the shortest path of the cell body, and the MreB/Fib association stabilize the copolymer structure. The fibril transmits the cumulative force generated by MreBs and itself to the membrane to form and propagate the kink along the cell helix. The role of the other MreBs (B2, B3 and B4) remains to be elucidated. As discussed above, it is plausible that MreB4 also constitutes the cytoskeleton ribbon, and that functions of MreB1 and MreB4 are overlapping. This study opens up new perspectives, in particular for understanding the swimming mechanism in *Spiroplasma*. Now that the involvement of MreBs in motility has been demonstrated, it appears essential to understand how the expression of MreBs is regulated during the cell cycle in order to distinguish their functions associated with division, from those linked to the maintenance of helicity and motility. Further studies will also be required to elucidate the mechanism of force generation by the cytoskeleton components and its transmission to the membrane, as well as to determine whether the tapered shape of the *Spiroplasma* end allows the propulsion of the cell in liquid medium.

## Materials & Methods

### Bacterial strains and culture conditions

*Escherichia coli* (NEB® 10-beta Electrocompetent *E. coli* or NEB® 5-alpha Competent *E. coli* High Efficiency) served as host strains for cloning experiments and plasmid propagation. Plasmid-transformed *E. coli* cells were grown at 37°C in Luria-Bertani (LB) broth or on LB agar supplemented with ampicillin at 100µg/ml and tetracycline 5 µg/mL.

The restriction free *M. capricolum* subsp. *capricolum* strain California KidT (*Mcap*) was used in this study. This strain was obtained by inactivation of the CCATC-restriction enzyme in the wild-type strain (ATCC 27343) (Lartigue et al., 2009). *Mcap* as well as its derivatives were grown at 37°C in SP5 medium (Tully et al., 1977) supplemented with tetracycline 5 µg/mL. One passage corresponds to the transfer of 10 μL of mycoplasma culture (at pH∼6.5) to 1 mL of mycoplasma medium (1/100 dilution) followed by an incubation period of ≥ 24h (depending on the constructions) at 37 °C. SP5 medium containing 0.8% noble agar was used to grow colonies of *Mcap* and its derivatives.

*S. citri* strain GII-3 was originally isolated from its leafhopper vector *C. haematoceps* captured in Morocco (Vignault et al., 1981). Spiroplasmas were cultivated at 32°C in SP4 medium. SP4 containing 0.8% noble agar was used to grow colonies of *S. citri*.

### Plasmid construction

Seven plasmids were built during this study. All derive from the transposon based-plasmid pMT85-PStetM (4.73 kbp) (Dordet-Frisoni et al. 2014; Zimmerman and Herrmann 2005; Aboklaish et al. 2014) which harbours the *tet*(M) gene from transposon Tn916 under the control of the spiralin promoter (PS).

Five plasmids were constructed using the NEBuilder® HiFi DNA Assembly Cloning Kit (Table S3 and Figures S1, S3, S4, S5). Depending on the assemblies, two to four overlapping DNA fragments were PCR amplified (Advantage HF 2 PCR Kit from Clontech) using primers described Table S4, purified and combined at 50°C according to the manufacturer’s instructions. DNA cassettes were designed to contain ∼40bp overlaps. One DNA cassette corresponds to the whole transposon based-plasmid pMT85-PStetM. Others correspond to different regions of *S. citri* GII-3 genome: the *fibril* gene (SPICI12_006), the *mreB1* gene (SPICI13_009) and the locus composed of *mreB2, mreB3*, a hypothetical protein encoding gene (HP), *mreB4* and *mreB5* (SPICI01A_045 to SPICI01A_049). *S. citri* DNA cassettes were amplified so that native promoters were conserved.

The two last plasmids (pMT85PStetM-mreB1 and pMT85PStetM-*mreB5*), which derived from plasmids pMT85PStetM-*mreB1-fib* and pMT85PStetM-*mreB5-fib* respectively, were build using the Q5® Site-Directed Mutagenesis Kit Protocol (Table S3).

All the oligonucleotides for plasmids constructions were supplied by Eurogentec and are described in Table S4.Prior to being used for transformation into *Mcap*, the purified plasmids were verified by restriction analyses and sequencing.

### Mycoplasma transformation and screening

*Mcap* was grown in SOB+ medium at 30°C and transformed using the established 5% polyethylene glycol (PEG) mediated protocol (Labroussaa et al., 2016). Transformations were conducted using 5 to 10 µg of plasmids and transformants were grown on selective medium SP5 plus tetracycline 5□µg/mL (SP5 tet5) for 7 to 21 days depending on the construction.

Colonies obtained on selective plates were picked and transferred into 1□mL of SP5 tet5 liquid medium and incubated at 37□°C. After three passages, 200 µL were used for DNA extraction and PCR analysis and 800µL were stored at −80□°C.

Transformants genomic DNA was extracted with the NucleoSpin® Tissue kit (Macherey-Nagel, Düren, Germany) and further analyzed by (i) PCR to verify the presence of the *tet*(M) and *S. citri* genes (*fibril, mreb1, mreb2-5*) and (ii) direct sequencing to localize the transposon insertion site (see below).

Primers used for PCR, direct sequencing and localization of insertions into *Mcap* genomes are summarized Table S4.

### Determination of the transposon insertion site by Single-Primer PCR

Transposon insertion sites were determined by single-primer PCR. The 25□µL final reaction volume contained 1X PCR Buffer (NEB), 3□mM MgCl2, 1□µM of primer SPP2-pMT85-TetM, 0.2□mM dNTPs, 0.5U of Taq NEB polymerase (NEB), and 2.5□µL of the transformant DNA. The PCR amplification cycle was performed as previously described (Dordet Frisoni et al., 2013).

Transposon insertion sites were determined by Sanger sequencing of the PCR products with the nested primer MT85-1 (Table S4).

### Dark-field microscopy, cell length measurements

One volume of cultures of exponentially growing *Mcap* in SP5 tet5 and of *S. citri* in SP4 (pH 6.9) was diluted in two volumes of fresh medium. Bacterial suspensions were prepared between two microscope slides, with a liquid thickness of 15 µm. The morphology of *Mcap* transformants growing in SP5 media was observed using an Eclipse Ni (Nikon) microscope working in reflection and equipped with a dark field condenser. The Nikon oil immersion microscope objective was a 60X with a numerical aperture (N. A.) of 0.80. Pictures were taken with a camera Iris 9™ Scientific CMOS (2960 × 2960 pixels). Videos were recorded at the maximal frame rate, 10 to 30 frames per second (fps) using the software NIS-Elements Br (Nikon). *Mycoplasma* cell length and helicity parameters were measured from isolated frames using the same software. Helical pitch measurements excluded non-helical cells. The data are expressed as the mean ± standard deviation (SD). Statistical significance was estimated by the unpaired t test, two tailed. Time-lapse images were isolated from the video recordings.

### Cryo Electron Microscopy

Lacey carbon formvar 300 mesh copper grids were used. They were first submitted to a standard glow discharged procedure (3 mbar, 3 mA for 40 s). Plunge freezing was realized using the EM-GP apparatus (Leica). Four microliters of the sample was deposited on the grid and immediately blotted for 2 s with a Whatmann paper grade 5 before plunging into a liquid ethane bath cooled with liquid nitrogen (−184 °C). The settings of the chamber were fixed at 80% humidity and 20 °C. Specimen were used non-diluted in culture medium. They were observed at −178 °C using a cryo holder (626, Gatan), with a ThermoFisher FEI Tecnai F20 electron microscope operating at 200 kV under low-dose conditions. Images were acquired with an Eagle 4k x 4k camera (ThermoFisher FEI) and processed in ImageJ.

### Proteomics

*Mycoplasma* and *Spiroplasma* cells were harvested by centrifugation at 10,000 g, 20 min, before being washed twice with Dulbecco’s phosphate-buffered saline (Eurobio) (mycoplasmas), or with a solution containing 8 mM Hepes and 280 mM sucrose (spiroplasmas). Protein concentration was determined using the DC Protein Assay (Bio-Rad). Fifteen micrograms of proteins were mixed with SDS loading buffer (75 mM Tris-HCl, pH 6.8, 50 mM DTT, 2% SDS, 0.02% bromophenol blue, 8.5% glycerol), then loaded onto a SDS-PAGE stacking gel (7%). A short electrophoresis was performed (10 mA, 45 min and 20mA, 2h) in order to concentrate proteins. After migration, gels were stained with Coomassie Blue and destained (50% ethanol, 10% acetic acid, 40% deionized water). The revealed protein band from each fraction was excised, washed with water, and then immersed in a reductive solution (5 mM DTT). Cysteines were irreversibly alkylated with 20 mM iodoacetamide in the dark. Following washing and drying steps, gel bands were submitted to protein digestion with trypsin added to a final protease to protein ratio of 1:25, for 3 hours at 37 °C, in ammonium bicarbonate buffer (10 mM and pH 8). Peptides were extracted with 50% CH_3_CN, followed by 0.1% TFA, and finally 100% CH_3_CN. The collected samples were then dried. For each clone, three technical replicates were performed. Tandem Mass Spectrometry. Peptides were then analyzed by mass spectrometry. All experiments were performed on a LTQ-Orbitrap Elite (Thermo Scientific) coupled to an Easy-nLC II system (Thermo Scientific). One microliter of sample (200 ng) was injected onto an enrichment column (C18 Acclaim PepMap100, Thermo Scientific). The separation was performed with an analytical column needle (NTCC-360/internal diameter: 100 μm; particle size: 5 μm; length: 153 mm, NikkyoTechnos, Tokyo, Japan). The mobile phase consisted of H_2_O/0.1% formic acid (FA) (buffer A) and CH_3_CN/FA 0.1% (buffer B). Tryptic peptides were eluted at a flow rate of 300 nL/min using a three-step linear gradient: from 2 to 40% B over 76 min, from 40 to 100% B in 4 min and 10 min at 100% B. The mass spectrometer was operated in positive ionization mode with capillary voltage and source temperature set at 1.8 kV and 275 °C, respectively. The samples were analyzed using CID (collision induced dissociation) method. The first scan (MS spectra) was recorded in the Orbitrap analyzer (*r* = 60,000) with the mass range *m*/*z* 400–1800. Then, the 20 most intense ions were selected for tandem mass spectrometry (MS^2^) experiments. Singly charged species were excluded for MS^2^ experiments. Dynamic exclusion of already fragmented precursor ions was applied for 30 s, with a repeat count of 2, a repeat duration of 30 s and an exclusion mass width of ±5 ppm. Fragmentation occurred in the linear ion trap analyzer with collision energy of 35. All measurements in the Orbitrap analyzer were performed with on-the-fly internal recalibration (lock mass) at *m*/*z* 445.12002 (polydimethylcyclosiloxane). Resulting raw files were analyzed with Proteome Discoverer 1.4.1.14 (Thermo Scientific). Database search was performed with Mascot (version 2.8.0) algorithm against “GCF_000012765.1_ASM1276v1_protein” (801 entries) retrieved from NCBI and containing the protein sequences encoded in *M. capricolum* subsp. *capricolum* genome, and a multifasta containing *S. citri* GII-3 and plasmid-encoded protein sequences. The following search parameters were used: trypsin was specified as the enzyme allowing for one mis-cleavage; carbamidomethyl (C) and oxidation (M) were specified as variable modifications; precursor mass range was set between 350 and 5000 Da, with a precursor mass tolerance and a fragment ion tolerance of 5 ppm and 0.3 Da respectively. Peptide validation was performed using Percolator algorithm (Käll et al., 2007) and only « high confidence » peptides were retained corresponding to 1% false discovery rate. A minimum of 2 PSMs and 2 unique peptides were required to consider protein identification.

## Supporting information

Table S4 (Primers)

## Acknowledgements

We thank Nicolas Martin, Jean-Christophe Baret (Univ. Bordeaux, CNRS-UMR5031, Bordeaux, France), and the ‘Frontiers of Life’ network (Univ. Bordeaux, France) for fruitful discussions. We also thank Sebastian Lillo and Bruce Richard for technical help. BL doctoral fellowship was granted by Univ. Bordeaux through the ‘Interdisciplinary projects’ call.

## Author contributions

Conceptualization and supervision: LB, CL; Data production: LB, CL, BL, FR, MH, MD, JH; Data analyses: LB, CL, MH, JH, MD, OL; Writing-original draft: LB, CL, AB; Writing-review and editing: All authors; Funding acquisition: LB, CL, JPD, AB.

