## Supplemental Figures S1-S5 and Tables S1-S3 for "Tuning spherical cells into kinking helices in wall-less bacteria"

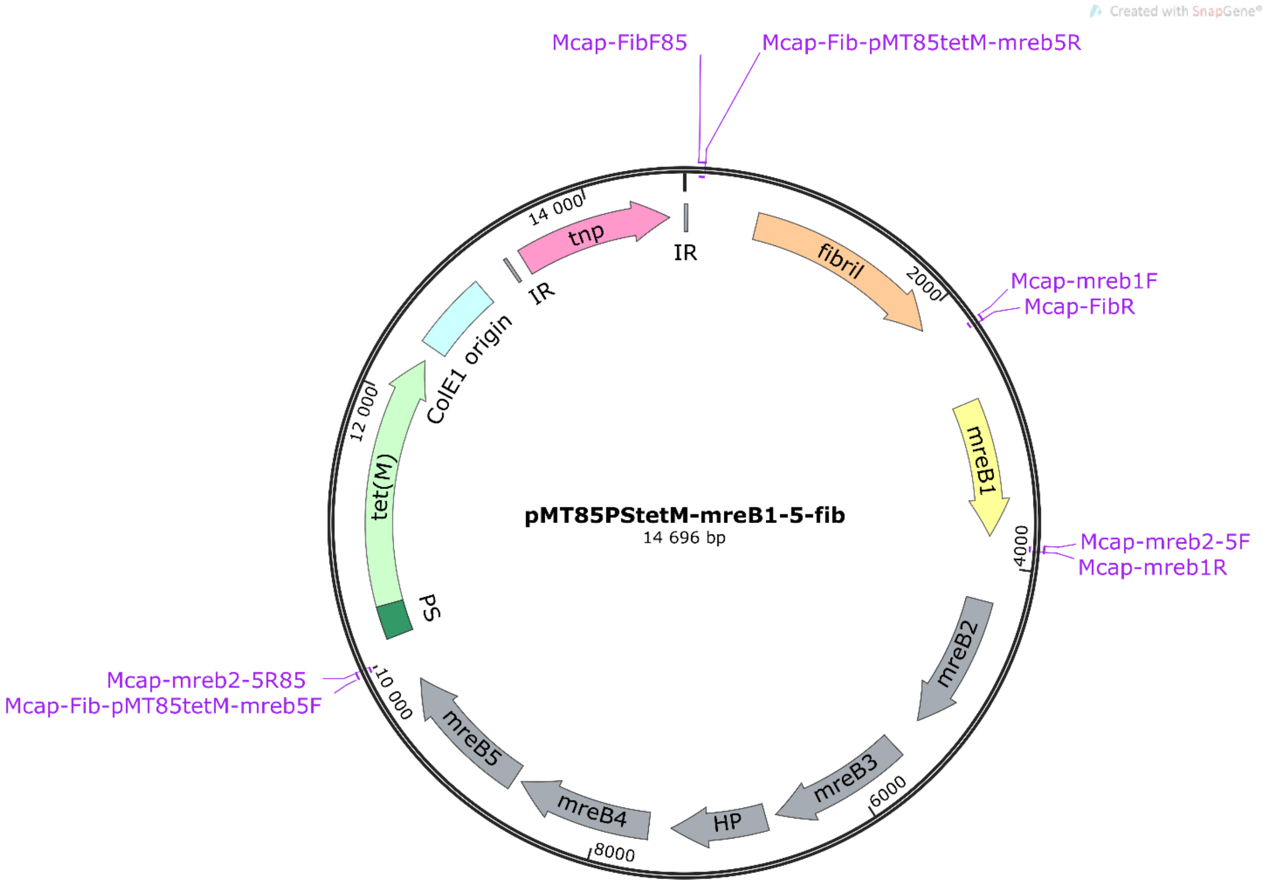

**Figure S1**. **Map of the pMT85PStetM-mreB1-5-fib plasmid.** This plasmid derives from the transposon based-plasmid pMT85-PStetM (4.82 kbp) which harbors the *tet*(M) gene (from *tn916*) under the control of the spiralin promoter (PS), a ColE1 origin and a transposase encoding gene (tnp) flancked by two inverted repeats (IR) from *tn4001* transposon. This plasmid contains 7 others genes (*fibril*, *mreB1*, *mreB2*, *mreB3*, an hypothetical protein encoding gene (HP), *mreB4* and *mreB5*) under the control of their native promoters. Primers used to assemble the plasmid are indicated in pink.

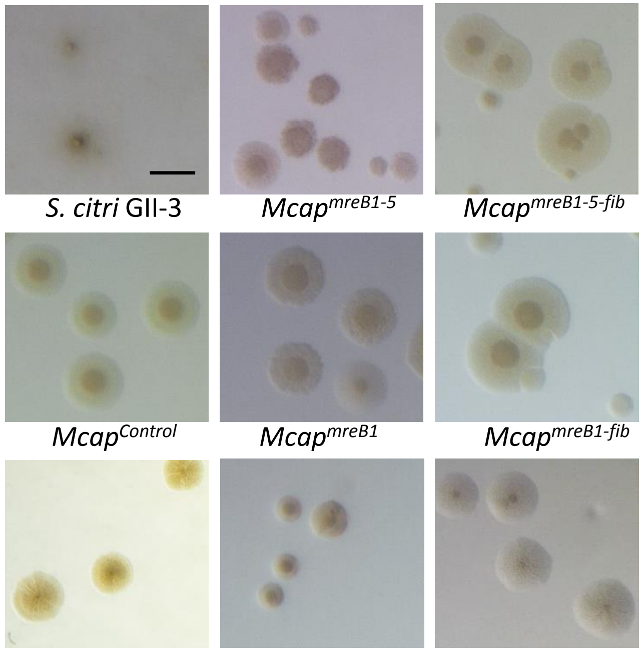

**Figure S2**. **Light microscopy images of colonies of *S. citri* GII-3 cells and of *Mcap* transformed with different *mreB* and *fib* gene combinations.** *Mcap*$^{control}$ corresponds to *Mcap* transformed with the pMT85-PStetM vector without any additional gene. Typical diffuse colonies were observed for *S. citri* with satellite colonies, and *Mcap*$^{control}$ produced fried-egg colonies. *Mcap* transformants did not form satellite colonies, and those having *mreB5* gene produced darker and smaller colonies. Irregular contour and/or granular aspect were observed for most *Mcap* transformants. Scale bar: 200 µm.

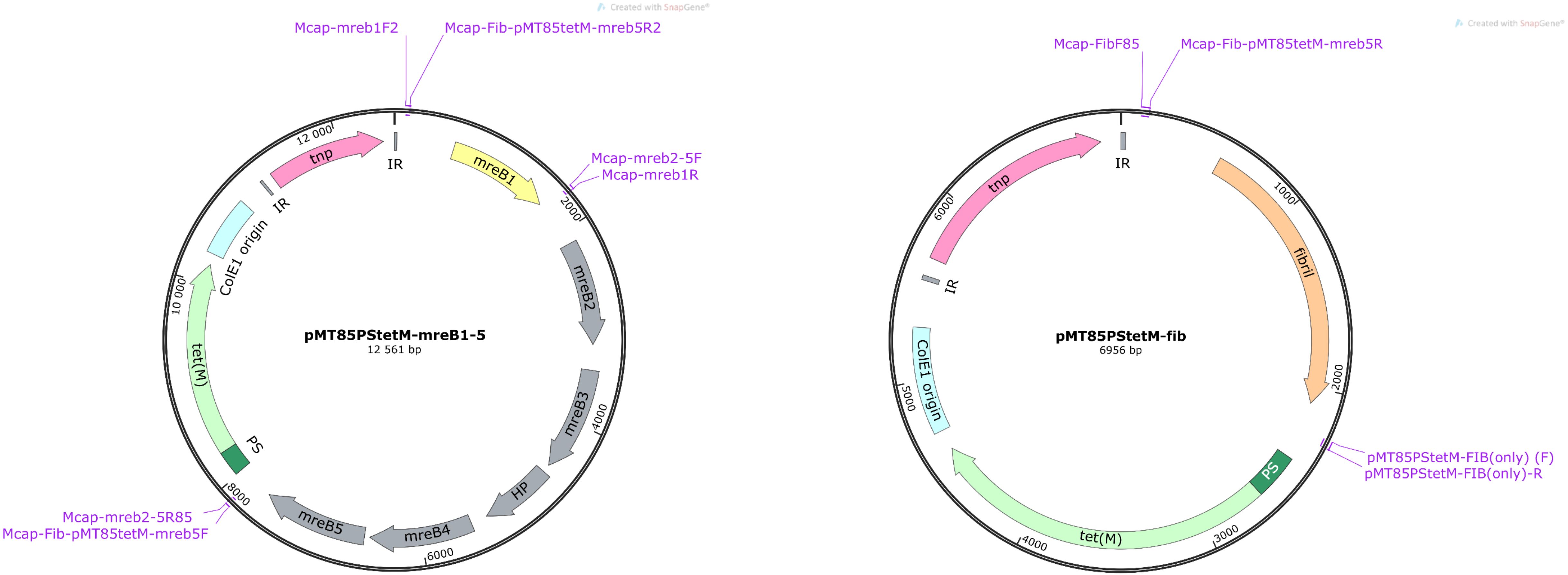

**Figure S3**. **Maps of the pMT85PStetM-mreB1-5 and pMT85PStetM-fib plasmids.** Both plasmids derive from the transposon based-plasmid pMT85-PStetM (4.82 kbp) which harbors the *tet*(M) gene (from *tn916*) under the control of the spiralin promoter (PS), a ColE1 origin and a transposase encoding gene (tnp) flancked by two inverted repeats (IR) from *tn4001* transposon. The pMT85PStetM-mreB1-5 plasmid contains 6 others genes (*mreB1*, *mreB2*, *mreB3*, an hypothetical protein encoding gene (HP), *mreB4* and *mreB5*) under the control of their native promoters while pMT85PStetM-fib plasmid harbors solely the *fibril* encoding gene under the control of its native promoter. Primers used to assemble both plasmids are indicated in pink.

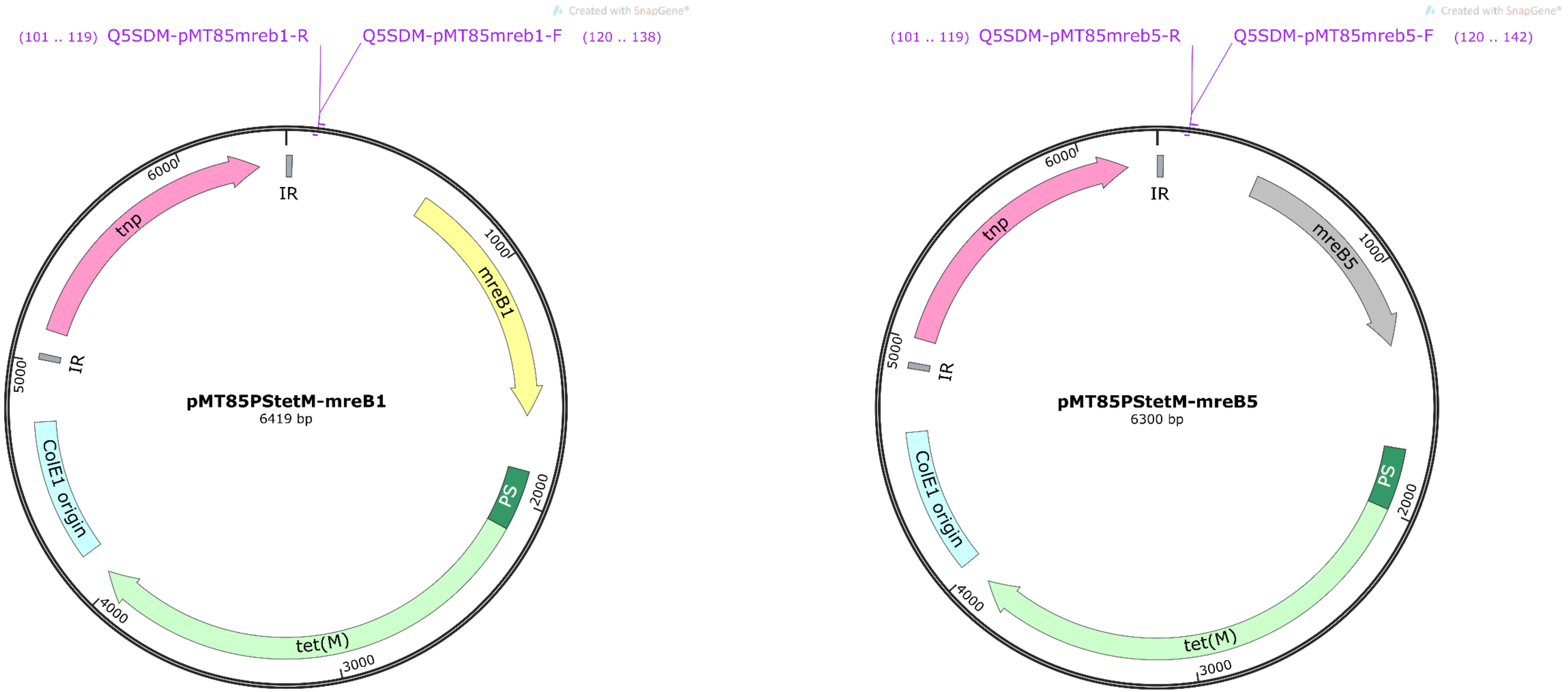

**Figure S4. Maps of the pMT85PStetM-mreB1 and pMT85PStetM-mreB5 plasmids.** Both plasmids derive from the transposon based-plasmid pMT85-PStetM (4.82 kbp) which harbors the *tet*(M) gene (from *tn916*) under the control of the spiralin promoter (PS), a ColE1 origin and a transposase encoding gene (tnp) flancked by two inverted repeats (IR) from *tn4001* transposon. The pMT85PStetM-mreB1 and pMT85PStetM-mreB5 contain the *mreB1* and *mreB5* encoding genes respectively, under the control of their native promoters. Primers used to assemble both plasmids are indicated in pink.

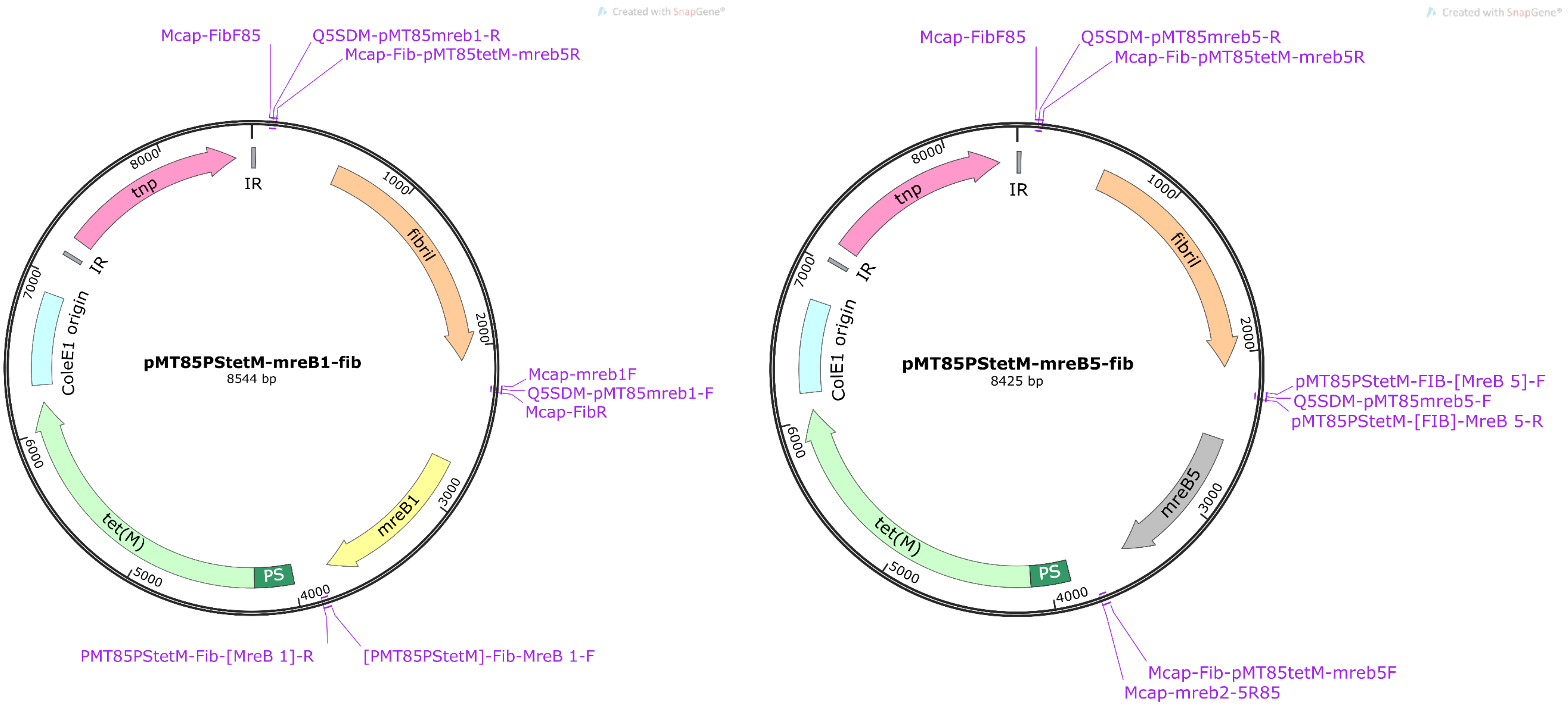

**Figure S5**. **Maps of the pMT85PStetM-mreB1-fib and pMT85PStetM-mreB5-fib plasmids.** Both plasmids derive from the transposon based-plasmid pMT85-PStetM (4.82 kbp) which harbors the *tet*(M) gene (from *tn916*) under the control of the spiralin promoter (PS), a ColE1 origin and a transposase encoding gene (tnp) flancked by two inverted repeats (IR) from *tn4001* transposon. The pMT85PStetM-mreB1-fib harbors the *mreB1* and *fib* encoding genes under the control of their native promoters while the pMT85PStetM-mreB5 harbors the *mreb5* and *fib* encoding genes under the control of their native promoters. Primers used to assemble both plasmids are indicated in pink.

**Table S1. Localisation of the added genes in the *Mcap* genome**

| Plasmids | Clone number | Integration site | Position in the *Mcap* genome (nt) |
|---|---|---|---|
| pMT85-PStetM | 2.1 | Mcap0559 - TraG/TraD | 690/2001 |
| pMT85-PStetM | 5.1 | Mcap0466 - Endopeptidase O | 236/1893 |
| pMT85PStetM-mreB1-5-fib | 8.7 | Mcap0222 - Threonyl tRNA synthetase | 859/1160 |
| pMT85PStetM-mreB1-5-fib | 32.1 | Mcap0846 - Lipoprotein | 1031/2091 |
| pMT85PStetM-mreB1-5 | 7.5 | Complete integration of the plasmid in the *Mcap* genome | NA |
| pMT85-PStetM-fib | 3.1 | Mcap0086 - Lipoprotein | 972/2091 |
| pMT85-PStetM-fib | 3.2 | Intergenic region between Mcap0791 (HP) and Mcap0792 (DNA Topoisomerase) | 913795/1010023 |
| pMT85-PStetM-fib | 8.4 | Mcap0791 - HP | 157/876 |
| pMT85-PStetM-fib | 3.22 | Mcap0460 - 6 Phospo gluconate dehydrogenase | 278/900 |
| pMT85-PStetM-mreB5 | 12.1 | Complete integration of the plasmid in the *Mcap* genome | NA |
| pMT85-PStetM-mreB5 | 24.3 | Complete integration of the plasmid in the *Mcap* genome | NA |
| pMT85-PStetM-mreB5-fib | 17.1 | Complete integration of the plasmid in the *Mcap* genome | NA |
| pMT85-PStetM-mreB5-fib | 17.3 | Mcap0362 - HP | 1900/2508 |
| pMT85-PStetM-mreB1 | 8.1 | Complete integration of the plasmid in the *Mcap* genome | NA |
| pMT85-PStetM-mreB1 | 7.1 | Complete integration of the plasmid in the *Mcap* genome | NA |
| pMT85-PStetM-mreB1-fib | 13.1 | Complete integration of the plasmid in the *Mcap* genome | NA |

NA: not applicable. Primers used during SPP allowed concluding that the entire transposon-based plasmid integrated the *Mcap* genome in these clones. However, they do not allow determining the site insertion of the transposon-based plasmid.

**Table S2.** ***Mcap*** **transformation efficiency with the plasmids built during this study and harboring different combination of** ***mreB*** **and** ***fibril*** **encoding gene(s)**

| **Plasmid** | **Transformation efficiencies* (Number tfs/mL/µg of plasmids)** |
|---|---|
| pMT85-PStetM | 3,15E-08 |
| pMT85PStetM-mreB1-5-fib | 1,85E-10 |
| pMT85PStetM-mreB1-5 | 7,21E-11 |
| pMT85-PStetM-fib | 1,85E-08 |
| pMT85-PStetM-mreB5 | 2,44E-11 |
| pMT85-PStetM-mreB5-fib | 2,00E-10 |
| pMT85-PStetM-mreB1 | 1,20E-10 |
| pMT85-PStetM-mreB1-fib | 1,70E-10 |

**Calculated from at least 3 independent experiments*

**Table S3. DNA cassettes used for plasmids constructions**

| **Plasmids Names** | **Number of DNA cassettes used for the Gibson assembly reaction** | **Cassette "pMT85PStetM"** (~4,862bp) | **Cassette "*fibril*"** SPICI12_006 (~2,175bp) | **Cassette "*mreB1*"** (SPICI13_009) (~1,637bp) | **Cassette "*mreB2-3-HP-4-5*"** (SPICI01A_045 to 049) (~6,185bp) |
|---|---|---|---|---|---|
| pMT85PStetM-mreB1-5-fib | 4 | + | + | + | + |
| pMT85PStetM-mreB1-5 | 3 | + | - | + | + |
| pMT85PStetM-fib | 2 | + | + | - | - |
| pMT85PStetM-mreB1-fib | 3 | + | + | + | - |
| pMT85PStetM-mreB5-fib | 3 | + | + | - | + *(mreB5 only)* |
| pMT85PStetM-mreB1 | Plasmid that derived from pMT85PStetM-mreB1-fib and built using the Q5® Site-Directed Mutagenesis Kit Protocol. | | | | |
| pMT85PStetM-mreB5 | Plasmid that derived from pMT85PStetM-mreB5-fib and built using the Q5® Site-Directed Mutagenesis Kit Protocol. | | | | |

**Table S4. → separate PDF file with primers**
