## Supplementary material for "Tuning spherical cells into kinking helices in wall-less bacteria": Table S4 (Primers)

### PCR primer pairs used to build plasmid pMT85PStetM-mreB1-5-fib

| Target region or template | Primers'names | Sequence (5'--> 3') | Amplicon size |
|---|---|---|---|
| Transposon based-plasmid pMT85-PStetM | Mcap-Fib-pMT85tetM-mreB5F | ccacggtatggatcaggagt cagctatgaccttgattacgg | 4,862 bp |
| | Mcap-Fib-pMT85tetM-mreB5R | cgatccaactattcaagcacc gtataaagtgtaaagcctgg | |
| *fibril* *(Spiroplasma citri GII3)* | Mcap-FibF85 | ccaggctttacactttatac ggtgcttgaatagttggatcg | 2,175 bp |
| | Mcap-FibR | ggatcttcatctggaatacc gttactatctccttctgctg | |
| *mreB1* *(Spiroplasma citri GII3)* | Mcap-mreB1F | cagcagaaggagatagtaac ggtattccagatgaagatcc | 1,637 bp |
| | Mcap-mreB1R | Tggcgtggctgaagcatctg catagttgctctatgctgacc | |
| *locus mreB2-3-HP-4-5* *(Spiroplasma citri GII3)* | Mcap-mreB2-5F | ggtcagcatagagcaactatg cagatgcttcagccacgcca | 6,185 bp |
|  | Mcap-mreB2-5R85 | ccgtaatcaaggtcatagctg actcctgatccataccgtgg |  |

### PCR primer pairs used to build plasmid pMT85PStetM-mreB1-5

| Target region or template | Primers'names | Sequence (5'--> 3') | Amplicon size |
|---|---|---|---|
| Transposon based-plasmid pMT85-PStetM | Mcap-Fib-pMT85tetM-mreB5F | ccacggtatggatcaggagt cagctatgaccttgattacgg | 4,862 bp |
| | Mcap-Fib-pMT85tetM-mreB5R2 | ggatcttcatctggaatacc gtataaagtgtaaagcctgg | |
| *mreB1* *(Spiroplasma citri GII3)* | Mcap-mreB1F2 | ccaggctttacactttatac ggtattccagatgaagatcc | 1,637 bp |
| | Mcap-mreB1R | Tggcgtggctgaagcatctg catagttgctctatgctgacc | |
| *locus mreB2-3-HP-4-5* *(Spiroplasma citri GII3)* | Mcap-mreB2-5F | ggtcagcatagagcaactatg cagatgcttcagccacgcca | 6,185 bp |
|  | Mcap-mreB2-5R85 | ccgtaatcaaggtcatagctg actcctgatccataccgtgg |  |

### PCR primer pairs used to build plasmid pMT85PStetM-fib

| Target region or template | Primers'names | Sequence (5'--> 3') | Amplicon size |
|---|---|---|---|
| Transposon based-plasmid pMT85-PStetM | pMT85PStetM-FIB(only) (F) | cagcagaaggagatagtaac cagctatgaccttgattacgg | 4,862 bp |
| | Mcap-Fib-pMT85tetM-mreB5R | cgatccaactattcaagcacc gtataaagtgtaaagcctgg | |
| *fibril* *(Spiroplasma citri GII3)* | Mcap-FibF85 | ccaggctttacactttatac ggtgcttgaatagttggatcg | 2,175 bp |
| | pMT85PStetM-FIB(only)-R | ccgtaatcaaggtcatagctg gttactatctccttctgctg | |

### PCR primer pairs used to build plasmid pMT85PStetM-mreB1-fib

| Target region or template | Primers'names | Sequence (5'--> 3') | Amplicon size |
|---|---|---|---|
| Transposon based-plasmid pMT85-PStetM | [PMT85PStetM]-Fib-MreB 1-F | ggtcagcatagagcaactatg cagctatgaccttgattacgg | 4,854 bp |
| | Mcap-Fib-pMT85tetM-mreB5R | cgatccaactattcaagcacc gtataaagtgtaaagcctgg | |
| *fibril* *(Spiroplasma citri GII3)* | Mcap-FibF85 | ccaggctttacactttatac ggtgcttgaatagttggatcg | 2,175 bp |
| | Mcap-FibR | ggatcttcatctggaatacc gttactatctccttctgctg | |
| *mreB1* *(Spiroplasma citri GII3)* | Mcap-mreB1F | cagcagaaggagatagtaac ggtattccagatgaagatcc | 1,637 bp |
| | PMT85PStetM-Fib-[MreB 1]-R | ccgtaatcaaggtcatagctg catagttgctctatgctgacc | |

### PCR primer pairs used to build plasmid pMT85PStetM-mreB5-fib

| Target region or template | Primers'names | Sequence (5'--> 3') | Amplicon size |
|---|---|---|---|
| Transposon based-plasmid pMT85-PStetM | Mcap-Fib-pMT85tetM-mreB5F | ccacggtatggatcaggagt cagctatgaccttgattacgg | 4,862 bp |
| | Mcap-Fib-pMT85tetM-mreB5R | cgatccaactattcaagcacc gtataaagtgtaaagcctgg | |
| *fibril* *(Spiroplasma citri GII3)* | Mcap-FibF85 | ccaggctttacactttatac ggtgcttgaatagttggatcg | 2,175 bp |
| | pMT85PStetM-[FIB]-MreB 5-R | ggtaattgtaatgtatctgt gttactatctccttctgctg | |
| *mreB5* *(Spiroplasma citri GII3)* | pMT85PStetM-FIB-[MreB 5]-F | cagcagaaggagatagtaac acagatacattacaattacc | 1,510 bp |
|  | Mcap-mreB2-5R85 | ccgtaatcaaggtcatagctg actcctgatccataccgtgg |  |

### PCR primer pairs used to build plasmid pMT85PStetM-mreB1

| Target region or template | Primers'names | Sequence (5'--> 3') | Amplicon size |
|---|---|---|---|
| pMT85PStetM-mreB1-fib | Q5SDM-pMT85mreB1-F | ggtattccagatgaagatc | 6,409 bp |
| | Q5SDM-pMT85mreB1-R | gtataaagtgtaaagcctg | |

### PCR primer pairs used to build plasmid pMT85PStetM-mreB5

| Target region or template | Primers'names | Sequence (5'--> 3') | Amplicon size |
|---|---|---|---|
| pMT85PStetM-mreB5-fib | Q5SDM-pMT85mreB5-F | acagatacattacaattaccatc | 6,290 bp |
| | Q5SDM-pMT85mreB5-R | gtataaagtgtaaagcctg | |

### PCR primer pairs used to verify the presence of foreign gene(s)

| Amplification | Name | Sequence 5'--> 3' | Amplicon size |
|---|---|---|---|
| *tet(M)* | Tet1 | ctgcaaaagatggcgtac | 535 bp |
| | Tet2 | cgtaaatgtagtactccac | |
| *mreB1* | Mcap-mreB1F | cagcagaaggagatagtaac ggtattccagatgaagatcc | 1,637 bp |
| | Mcap- mreB1R | Tggcgtggctgaagcatctg catagttgctctatgctgacc | |
| *mreB2-5* | Mcap-mreB2-5F | ggtcagcatagagcaactatg cagatgcttcagccacgcca | 6,185 bp |
|  | Mcap- mreB2-5R85 | ccgtaatcaaggtcatagctg actcctgatccataccgtgg |  |
| *fibril* | Mcap-FibF85 | ccaggctttacactttatac ggtgcttgaatagttggatcg | 2,175 bp |
| | Mcap-FibR | ggatcttcatctggaatacc gttactatctccttctgctg | |
| *mreB1* *(in pMT85PStetM-mreB1-fib and pMT85PStetM-mreB1)* | Mcap-MreB1F | cagcagaaggagatagtaac ggtattccagatgaagatcc | 1,637 bp |
| | PMT85PStetM-Fib-[MreB 1]-R | ccgtaatcaaggtcatagctg catagttgctctatgctgacc | |
| *mreb5* *(in pMT85PStetM-mreB5-fib and pMT85PStetM-mreB5 )* | pMT85PStetM-FIB-[MreB 5]-F | cagcagaaggagatagtaac acagatacattacaattacc | 1,546 bp |
| | Mcap-MreB2-5R85 | ccgtaatcaaggtcatagctg actcctgatccataccgtgg | |

### PCR primers used for localization of the transposon by single priming PCR and sequencing

| Amplification | Name | Sequence 5'--> 3' | SPP Primers used for constructions |
|---|---|---|---|
| Mcap transformants | SPP-pMT85-TetM | ggtcatagctgtttcctgtg (for single priming PCR) | pMT85PStetM |
| Mcap transformants | SPP1-pMT85-fib | gatccaactattcaagcacc (for single priming PCR) | pMT85PStetM-fib<br>pMT85PStetM-mreB1-fib<br>pMT85PStetM-mreB5-fib<br>pMT85PStetM-mreB1-5-fib |
| Mcap transformants | SPP3-pMT85-mreB1 | actaattggatcttcatctgga (for single priming PCR) | pMT85PStetM-mreB1<br>pMT85PStetM-mreB1-5 |
| Mcap transformants | SPP4-pMT85-mreB5 | gatggtaattgtaatgtatctgt (for single priming PCR) | pMT85PStetM-mreB5 |
| Mcap transformants | MT85-1 | acagtaattgcgggtggatc (for sequencing) | Used for the sequencing step |
